# NMNAT2 in cortical glutamatergic neurons exerts both cell and non-cell autonomous influences to shape cortical development and to maintain neuronal health

**DOI:** 10.1101/2022.02.05.479195

**Authors:** ZhenXian Niou, Sen Yang, Anoosha Sri, Hugo Rodriquez, Jonathan Gilley, Michael P. Coleman, Hui-Chen Lu

## Abstract

Here we show that deleting NMNAT2 from cortical glutamatergic neurons (NMNAT2 cKO) results in progressive axonal loss, neuroinflammation, small hippocampi and enlarged ventricles. Interestingly, dramatic neuroinflammation responses were observed around the long-range axonal tracts of NMNAT2 cKO cortical neurons. In addition to the neurodegenerative-like phenotype, we also found the absence of whisker-representation patterns “barrels” in the primary somatosensory cortex of NMNAT2 cKO mice. These observations suggest that NMNAT2 is required in developing cortical circuits and in maintaining the health of cortical neurons. Unbiased transcriptomic analysis suggests that NMNAT2 loss in cortical neurons after axonal outgrowth phase upregulates mitochondria function while greatly reducing synaptogenesis pathways. Complete loss of Sarm1 function in NMNAT2 cKO mice restores barrel map formation and axonal integrity and abolishes the inflammatory response. Interestingly, reducing Sarm1 function in NMNAT2 cKO mice by deleting only one copy of Sarm1 restores barrel map formation but did not diminish the neurodegenerative-like phenotype. Only complete loss of Sarm1 prevents neurodegeneration and inflammatory responses.

## Introduction

Nicotinamide mononucleotide (NAD) is critical for energy metabolism and is involved in many pathways necessary for health and longevity (Bogan and Brenner, 2008; Chiarugi et al., 2012; Lautrup et al., 2019; Rajman et al., 2018; Verdin, 2015; Yoshino et al., 2018). Nicotinamide mononucleotide adenylyl transferases 1-3 (NMNAT1-3) are known as NAD synthesizing enzymes catalyzing the last step in the NAD salvage pathway in all living organisms (Ali et al., 2013a; Brazill et al., 2017; Conforti et al., 2014). In addition, we and others showed NMNATs can also function as molecular chaperones (Ali et al., 2016; Zhai et al., 2008). The overexpression of NMNATs offers potent neuroprotection in a variety of axonal injuries and neurodegenerative conditions (Ali et al., 2016; Gilley et al., 2013a; Gilley and Coleman, 2010; Ljungberg et al., 2012; Milde et al., 2013b). NMNAT2 is primarily expressed in differentiated neurons and is the most abundant NMNAT in mammalian brains (Berger et al., 2005; Raffaelli et al., 2002; Yan et al., 2010). NMNAT2 turns over quickly, with a half-life of less than 4 hrs *in vitro* (Gilley and Coleman, 2010; Milde et al., 2013a), significantly shorter than NMNAT1 and NMNAT3. Thus, its abundance is quickly reduced following axon injury, mitochondria insult or toxin exposure (Coleman and Hoke, 2020; Loreto et al., 2020; Yamagishi and Tessier-Lavigne, 2016). In addition to the labile nature of NMNAT2 protein, the abundance of NMNAT2 mRNA has also been shown to be reduced in the brains of Alzheimer’s disease (AD) (Ali et al., 2016), Parkinson’s disease (PD), Huntington’s disease (HD) patients and in the spinal cord of amyotrophic lateral sclerosis (ALS) patients (Ali et al., 2013b; Harlan et al., 2020).

Neurons often possess extensively arborized axonal processes that navigate over long distances to reach their targets (Winnubst et al., 2019). Axonopathy is often an early sign of neurodegenerative conditions (Fischer et al., 2004; Xiao et al., 2011). The importance of NMNAT2 in axonal health of peripheral neurons upon injury or stress has been primarily explored with peripheral nervous system and nerve injury models (Coleman and Hoke, 2020; Hughes et al., 2021; Simon and Watkins, 2018). These studies provide strong evidence supporting the importance of the NAD-synthesizing function of NMNAT2 in axonal health (Araki et al., 2004; Milde et al., 2013a; Yan et al., 2010). A sharp decline in axonal NAD levels together with the activation of Sarm1 (sterile alpha and TIR motif containing protein 1), a newly discovered NAD(P) glycohydrolase, were ascribed to axonal degeneration (Angeletti et al., 2022; Di Stefano et al., 2015; Gerdts et al., 2015). Global NMNAT2 knockout (KO) results in premature death at birth with a severe peripheral nerve defect and truncated axons in the optic nerve (Gilley et al., 2013a; Hicks et al., 2012). In NMNAT2 germline cKO mice, the neurite outgrowth phenotype can be rescued by overexpressing Wallerian degeneration slow (Wlds) chimeric protein, which provides exogenous nicotinamide mononucleotide adenylyl transferase activity to rescue the deficits caused by NMNAT2 insufficiency in the neurites (Gilley et al., 2013a).

The identification of Sarm1 as a key player in an axon-autonomous self-destruction program from a *Drosophila* genetic screen made clear that axonopathy is not due to passive wasting away of axons due to lack of nutrients from soma (Osterloh et al., 2012). Sarm1 activation triggers axon destruction (Essuman et al., 2017) while Sarm1 loss-of-function potently protects axons from wide range of insults (Coleman and Hoke, 2020; Figley and DiAntonio, 2020; Krauss et al., 2020), including injury-induced “Wallerian” degeneration and chemotherapy-induced peripheral neuropathy. Excitingly, SARM1 inactive variants and inhibitors can prevent axon degeneration (Bosanac et al., 2021; Essuman et al., 2017; Figley et al., 2021; Gould et al., 2021; Hughes et al., 2021). Despite being a member of the Myd88 immune adaptor protein family, SARM1 itself has no clear role in innate immunity. SARM1 catalyzes NAD^+^ hydrolysis (Figley and DiAntonio, 2020; Gerdts et al., 2015; Gerdts et al., 2016; Jiang et al., 2020) and alternatively possesses transglycosidation (base exchange) activity (Angeletti et al., 2022). These enzymatic activities are all activated by an increase in the ratio of nicotinamide mononucleotide (NMN) to NAD^+^, even without axonal injuries (Figley et al., 2021). SARM1 activation in axons is normally restrained by NMNAT2, which uses NMN and ATP to synthesize NAD (Garavaglia et al., 2002). Figley *et al*. (2021) showed that NMNAT2 reduction increases the ratio of NMN to NAD and activates SARM1 to trigger axon degeneration (Figley et al., 2021). This likely explains why SARM1 deletion was able to prevent the detrimental effects on survival and axon integrity caused by NMNAT2 loss of function (Di Stefano et al., 2015; Gilley et al., 2017).

Our previous studies showed that *nmnat2* mRNA levels in the prefrontal cortex of aged human subjects positively correlate to cognitive function and negatively correlate to AD pathology (Ali et al., 2016)). Despite the wealth of knowledge on the importance of NMNAT2 in peripheral nerves, the premature death of NMNAT2 KO mice has prevented the evaluation of the role(s) of NMNAT2 in the postnatal development of cortical neurons. Here, we specifically deleted NMNAT2 in post-mitotic glutamatergic neurons using a Cre-LoxP approach and characterized the anatomical phenotypes caused by NMNAT2 loss-of-function to explore the roles of NMNAT2 in cortical neurons.

## Material and Method

### Animals

The generation of NMNAT2^f/f^ mice carrying a Cre recombinase dependent gene trap cassette within the exon 1 of NMNAT2 gene have been described (Gilley et al., 2013a). Here we generated NMNAT2 conditional knockout (cKO) mice by crossing the NMNAT2^f/f^ mouse line with NEX-Cre mice (Goebbels et al., 2006) to delete NMNAT2 in cortical glutamatergic neurons after embryonic day 11.5 (E11.5). NMNAT2 cKO mice were smaller in size and exhibited an ataxia phenotype with wobbling movements. Thus, a 5gram DietGel 76A (Clear H_2_O, Westbrook, ME, USA) was given daily after weaning. Sarm1 KO (S^null^/S^null^) mice (Kim et al., 2007) were crossed with NMNAT2 cKO mice to generate cKO;S^null^/+ and cKO; S^null^/S^null^ transgenic mice. (The day on which the vaginal plug was detected was designated as E0.5. The first 24 h after birth was referred to as P0). Both male and female mice were used. All mice were housed under standard conditions with food and water provided ad libitum and maintained on a 12 hr dark/light cycle. Mice were treated in compliance with the guidelines of the U.S. Department of Health and Human Services and all procedures were approved by the Indiana University Bloomington Institutional Animal Care and Use Committee.

### Genotyping

Tissue lysates were prepared by immersing small pieces of mouse ears into the digestion buffer (50 mM KCl, 10 mM Tris–HCl, 0.1% Triton X-100, 0.1 mg/ml proteinase K, pH 9.0) for 3 h at 60°C. Lysates were then heated at 95°C for 20 min to inactivate proteinase K and then centrifuged at 16,000 rpm for 15 min. The supernatants were used as the DNA template for polymerase chain reactions (PCRs) using EconoTaq Plus Green 2X mater mix (Lucigen, Middleton, WI, USA). For NMNAT2 floxed allele, genotyping was conducted using the following primers: NM-A 5’-TGTTCTGAAACGTGACTCGCT-3’; NM-B 5’-GAGGCATAAACCCGGCATAGT-3’; NM-C 5’-AATCAGTCATAGACACTAGACAATCG-3’. The PCR products for WT and NMNAT2^f^ allele are 675 bp and 556 bp. The sequences for NEX-Cre genotyping primers are: NEX-F: 5’-GAG TCC TGG AAT CAG TCT TTT TC-3’; NEX-R: 5’-AGA ATG TGG AGT AGG GTG AC-3’; NCre-R: 5’-CCG CAT AAC CAG TGA AAC AG-3’. The PCR products for WT and Cre alleles are 770 bp and 520 bp. Sequences for Sarm^null^ (S^null^) genotyping primers are: Sarm1-common: 5’-GAAATGCATGGAGGGGTTG-3’; WT-R: 5’-CCACCAAACGTGTCCAATC-3’; Mut-R: 5’-TGT GGTTTCCAAATGTGTCAG-3’. The PCR products for WT and Sarm1 null alleles are 302 bp and 248 bp.

### Immunostaining

Mice were anesthetized and transcardially perfused with 4 % paraformaldehyde (PFA, Sigma, St. Louis, MO, USA) in phosphate buffered saline (PBS). The brains were harvested and post-fixed overnight in 4% PFA at 4°C. After rinsing with PBS, some brains were cryopreserved with 30% sucrose in PBS at 4°C for 1-2 days while some brains were kept in PBS. The cryo-preserved brains were mounted in OCT compound (Tissue-Tek O.C.T., Sakura, Torrance, CA, USA) on the cutting platform of a Leica Microtome SM-2000R or a cryotome (CM-1850, Leica) at −20°C and cut at 30-40 μm thickness in the coronal plane. Brains kept in PBS were sectioned with a Leica VT-1000 vibrating microtome (Leica Microsystems). Free-floating sections were used in all subsequent staining steps. The staining procedure was carried out at room temperature except for a few steps noted below. Sections were washed with PBS containing 0.3% Triton X-100 (0.3% PBST) 5 min three times, permeabilized with 0.3% PBST for 20 min, incubated with blocking solution (3% Bovine Serum Albumin (BSA, Sigma A9418) in 0.3% PBST) for 2hr, and then incubated with the primary antibodies diluted in blocking solution for overnight at 4°C. The next day, sections were washed with 0.3% PBST for 5 min three times and incubated with secondary antibodies diluted in blocking solution at 4°C for 2hr. After incubation, samples were washed with 0.3 % PBST three times. Draq5 (1:10,000 dilution, Cell Signaling) or 4’,6-diamidino-2-phenylindole (DAPI, 5 μg/ml, Invitrogen) were added during the first wash step to visualize nuclei. Dako Mounting Medium was used to mount the brain sections.

### IdU Labeling

10 mg/kg 5-Iodo-2’-deoxyuridine (IdU, MP Biomedical, Irvine, CA, USA) was injected into pregnant dams at 14.5 days post gestation. Four days later the IdU labeled embryos were harvested from anesthetized dam’s uterus, fixed with 4% PFA in PBS at 4°C for 24 h and then cryopreserved for 24 h with 30% sucrose in 1xPBS at 4°C.

### Microscopy and image Processing

Single plane fluorescent images were taken with a Leica TCS SPE confocal microscope (Leica DM 2500) with 10x (0.3 N.A.) and 40x (0.75 N.A.) objective lens or with a Nikon A1R laser scanning confocal (Nikon A1) with 10x (0.5 N.A.) objective lens. DAPI/Draq5 immunofluorescence was used to match corresponding sections and identify the same anatomical landmark to image. A minimum of three sections were imaged per mouse brain, and selected anatomical regions were comparable across all brains. Bright field images were obtained by Zeiss SteREO Discovery.V8 Microscope (Carl Zeiss Microscopy). Figures were made with Adobe Photoshop CS6 and Illustrator CS6 and brightness/contrast, orientations, and background corrections were applied to better illustrate the staining patterns.

### Quantitative analysis of confocal images

Images were analyzed using Fiji (Image J with updated plug-ins). Densities of IdU-, Iba1-, or S100b-positive cells were determined by manually marking positive cells using the Image J cell count function. 0.36 mm^2^ (0.6×0.6) of the primary somatosensory cortex and 0.075 mm^2^ (0.5×0.15) hippocampal CA1 area were imaged to quantify cell densities. For IdU^+^ cell quantification in embryonic brains, the cortical plate region spanning from pial surface to ventricle was selected and divided into 0.01 mm^2^ bins (0.2×0.05 mm) to quantify cell densities. Cortical thickness of the primary somatosensory cortex was measured from the pia surface to the edge of the corpus callosum from both hemispheres with acquired brightfield images.

For all quantification of immunostaining outcomes images were acquired with identical imaging parameters chosen to avoid saturated signals in any sample. The thickness of corpus callosum was measured from the dorsal to ventral edges of NFM+ axons using coronal plane sections. For NFM+ axonal tracts passing through the dorsal striatum, coronal slices only from the middle rostral-caudal axis were selected for quantification. Areas of blood vessels were manually excluded. The vessel-subtracted dorsal striatal areas were then “Thresholded” with the top 8-11 % tail of total pixels. Next, “Analyze particles” was used to acquire percentage of pixels. To quantify GFAP+ astrocytes, 0.075 mm^2^ (0.5×0.15 mm) of the hippocampus CA1 and 0.25 mm^2^ (0.5×0.5 mm) of the dorsal striatum were used. GFAP images “Thresholded” with top 3.7-4.5 % tail of total pixel and then “Analyze particles” function was used to acquire pixels percentages.

VGluT2 and PSD95 immunoreactivity were quantified from 10x images acquired from samples collected and stained simultaneously. 0.09 mm^2^ (0.6*0.15 mm) inside the L4 S1 whisker-barrel or L5 region were selected to measure VGluT2 and PSD95 signals by using “Analyze -> Measure -> IntDen” to acquire pixel intensities. L4 measurements were subtracted with L5 intensity (background) to acquire the VGluT2 or PSD95 intensities.

All quantifications were done blind to genotype information. Both hemispheres of the same animals were used for quantification. Contours were drawn in specific regions using DAPI immunofluorescence as a guide, and only cells within each selected region were counted.

### NMNAT2 In Situ Hybridization

In situ hybridization on embryonic sections was done by the Baylor College of Medicine Advanced Technology Core Labs. The experiment was performed using a high-throughput automated platform. Cryostat sections of E14.5 embryos were placed on a standard microscope slide that was subsequently incorporated into a flow-through chamber. The chamber was then placed into a temperature-controlled rack and the required solutions for pre-hybridization, hybridization, and signal detection reactions were added to the flow-through chamber with an automated solvent delivery system. Details can be found in a previous publication (Yaylaoglu et al., 2005). The RNA probe was generated using the sequence from the Allen Brain Atlas data base. Images were acquired by Leica CCD camera with a motorized stage. Multiple images were collected from the same section. Individual images are stitched together to produce a mosaic representing the entire section.

### RNA-seq and Gene Ontology (GO) analysis

NMNAT2^f/f^ cortical neurons were plated at a density of 10*10^5^/well in PDL coated 12 well plate. 2MOI lentivirus expressing iCre recombinase was added to culture at DIV9 to knockout Nmnat2 (cKO group), while 2 MOI lentivirus expressing copGFP was added to the control group. At DIV12, total RNA was extracted from control or cKO group (6 biological replicates per group) by RNeasy Mini Kit (Qiagen, Qiagenstr, Hilden, Germany) and followed by on column DNase digestion according to the manufacturer’s instruction. mRNA sequencing was performed by Center for Medical Genomics, Indiana University School of Medicine. The concentration and quality of total RNA samples was first assessed using the Agilent 2100 Bioanalyzer. The RNA integrity number was higher than 9.4 for all samples. 100ng of RNA per sample as used to prepare a dual-indexed strand-specific cDNA library using KAPA mRNA HyperPrep Kits (KK8581, Roche), that enriched for polyadenylated mRNA. The resulting libraries were assessed for quantity and size distribution using Qubit and Agilent 2100 Bioanalyzer. 1.5 picomolar pooled libraries were sequenced with 75 bp single-end configuration on NextSeq500 (Illumina) using NextSeq 500/550 High Output Kit. A Phred quality score (Q score) was used to measure the quality of sequencing. More than 90% of the sequencing reads reached Q30 (99.9% base call accuracy). The sequencing data were first assessed using FastQC (Babraham Bioinformatics, Cambridge, UK) for quality control. Then all sequenced libraries were mapped to the mouse genome (UCSC mm10) using STAR RNA-seq aligner. Genes with read count per million (CPM) > 0.5 in more than 3 of the samples were retained. The data was normalized using TMM (trimmed mean of M values) method. Differential expression analysis was performed using edgeR. False discovery rate (FDR) was computed from p-values using the Benjamini-Hochberg procedure. Genes with FDR < 0.05 were considered significantly altered and further separated into upregulated and downregulated list based on its fold change. The gene ontology overrepresentation analysis was conducted separately for up and downregulated genes, using web-based Enrichr with 2021 GO Terms for Biological Process (Kuleshov et al., 2016; Xie et al., 2021). The ingenuity pathway (IPA) analysis for up and downregulated genes was also performed in parallel to visualize the canonical pathways enriched with differentially expressed genes.

## Results

### Deleting NMNAT2 in cortical glutamatergic neurons results in reduced survival rates and the survivors have reduced body weights and deformed brains

Using *in situ* hybridization with an anti-sense RNA probe specifically detecting *nmnat2* mRNA in embryonic day 14.5 (E14.5) brain sections, we found that nmnat2 mRNA expression was enriched in the cortical plate (CP) where post-mitotic neurons are located (Fig. 1A). The transgenic mouse line carrying a conditional allele of NMNAT2 (abbreviated as NMNAT2^f^ for simplicity) was generated by inserting a Cre recombinase dependent gene trap cassette within intron 1 of the NMNAT2 gene (Gilley et al., 2013b). To determine the requirement of NMNAT2 in glutamatergic neurons after they entered the post-mitotic phase, we disrupted NMNAT2 expression in cortical glutamatergic neurons after E11.5 by crossing NMNAT2^f/f^ mice with NEX-Cre transgenic mice (Goebbels et al., 2006).

**Fig 1.**
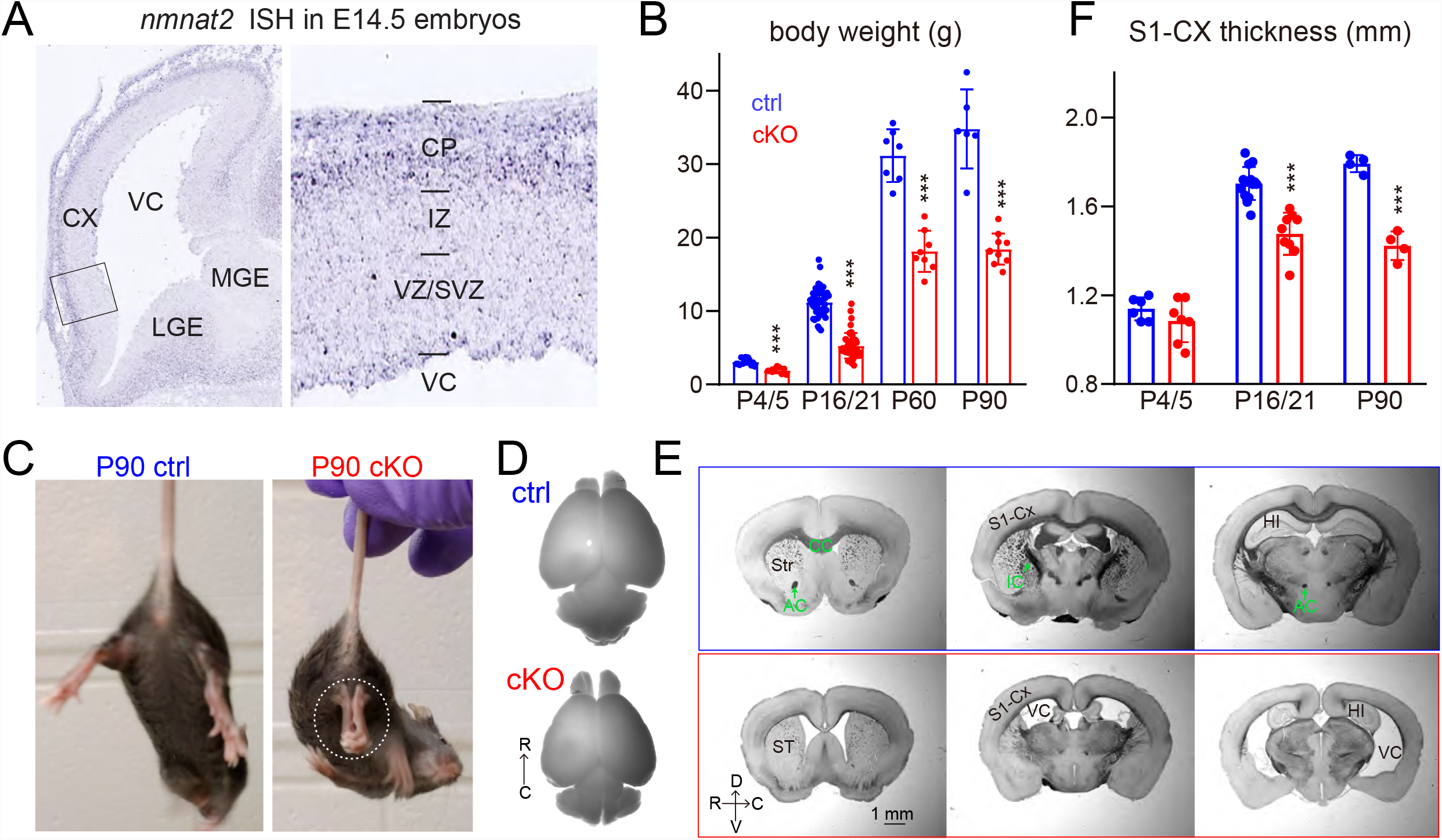
NMNAT2 expression and general characterization of NMNAT2 cKO mice. (**A**) In situ hybridization reveals *nmnat2* mRNAs (blue signals) in the E14.5 brain, mainly localized to the cortical plate (CP), the lateral ganglionic eminence (LGE) and the medial ganglionic eminence (MGE). (**B**) Summaries for body weights of control (ctrl) and NMNAT2-cKO mice at P4-P90 (P4/5 ctrl n=13, cKO n=9; P16/P21 ctrl n=43, cKO n=46; P60 ctrl n=7, cKO n=8, P90 ctrl n=6, cKO n=10). (**C**) Movie screen shots show a P90 cKO mouse but not a ctrl mouse exhibiting hindlimb clasping behavior (dashed white oval), a classic motor deficit observed in many neurodegenerative models. (**D, E**) Bright field images showing representative images for whole brains and coronal plane brain sections (rostral to caudal from left to right) from ctrl and cKO mice. In additional to a small brain size, cKO brains have enlarged ventricles and reduced cortical and hippocampi areas. (P4/5 ctrl n=3, cKO n=3; P16/21 ctrl n=9, cKO n=8; P90 ctrl n=3, cKO n=3) (**F**) Summaries for the thickness of the primary somatosensory (S1) cortex in ctrl and cKO mice at different ages (P4/5 Ctrl n=6, cKO n=7; P16/21 ctrl n=14, cKO n=9; P90 ctrl n=4, cKO n=4). Abbreviations: AC, anterior commissure; CX, cortex; CC, corpus callosum; HI, hippocampus; IC, internal capsule; IZ, intermediate zone; ST, striatum; SVZ, subventricular zone; VC, ventricle; VZ, ventricular zone. ***, p<0.001 by Student’s t-test.

To our surprise, despite the cortical-specific deletion of NMNAT2, only about 50% of the conditional NMNAT2 knockout (cKO) mice survived after birth (Sup-Fig. 1A) without obvious lethality at E18.5 (Sup-Fig. 1B). In surviving cKO mice, only ∼30% were female (7 out of 24 total), less than the 42.5% in control (17 out of 40 total mice). cKO mice weighed significantly less than their littermate controls from early postnatal ages to adulthood (Fig.1B). Around weaning age, NMNAT2 cKO mice exhibited evident hindlimb clasping (Fig. 1C), ataxia and forelimb circling phenotypes (data not shown; 10 out of 10 mice examined), reflecting motor behavior deficits, similarly observed in many neurodegenerative mouse models (Chou et al., 2008; Lieu et al., 2013; Wang et al., 2017). At the gross level, cKO brains were smaller than controls (Fig. 1D). We evaluated gross brain morphology of both cKO and littermate controls with coronal brain sections from rostral to caudal regions with bright field imaging. cKO brains displayed enlarged ventricles, smaller hippocampi, aberrant anterior commissures, a thinner corpus callosum and a thinner cortex (Fig. 1E). We quantitatively measured the thickness of the primary somatosensory (S1) cortex and found it was significantly reduced in cKO compared to their littermate controls at postnatal day 16-21 (P16/21) and P90 (Fig. 1F).

To investigate whether the smaller brains in cKO mice were caused by defective neuroprogenitor proliferations or radial migration occurring during embryonic stages, birth-dating labeling with IdU, an analog of thymidine integrating to DNA during S-phase of cell cycle, was conducted. Specifically, IdU was injected peritoneally to dams at E14.5 to label proliferating cells. The numbers and distributions of IdU+ in the cortical plate in cKO and littermate control brains were quantitatively examined. We found no difference for either IdU+ density or distribution from the ventricular/subventricular zone (VZ/SVZ) to the cortical plate (CP) between cKO and control brains. (Sup-Fig. 2). These data are consistent with the normal embryonic brain morphologies observed in NMNAT2 total KO mice (Hicks et al., 2012) and suggest that NMNAT2 is not required for the proliferation of neural progenitor cells and migration of post-mitotic neurons.

### NMNAT2 in cortical glutamatergic neurons is required for axonal health

Studies with the cortical explants prepared from NMNAT2 KO embryonic brains suggest the role of NMNAT2 in neurite outgrowth (Gilley et al., 2013a), while knocking-down NMNAT2 in mouse superior cervical ganglia (SCG) neurons led to Wallerian-like degeneration in axons (Gilley and Coleman, 2010). To examine the impacts of NMNAT2 loss on long-range axons originating from cortical glutamatergic neurons, immunostaining of neurofilament middle (NF-M), a cytoskeleton protein abundant in axons, was conducted with brain sections in the coronal plane prepared from P4/5, P16/21, and P90 cKO and littermate control brains (Fig. 2). We particularly focused on the corpus callosum, with cortical axons crossing to the contralateral hemisphere, and the striatum, where cortical axons project toward subcortical structures or the spinal cord. At P4/5, when the majority of cortical axons have finished midline crossing (Fame et al., 2011; Wang et al., 2007), we found that the thickness of cKO corpus callosum was similar to littermate controls (Fig. 2A, B). The abundance of axonal tracts projecting through cKO striatum was also similar to control striatum at P4/5 (Fig. 2C, D). But as the mice develop, they start to show a neurodegenerative phenotype at the second postnatal week. At P16/P21, both the thickness of the corpus callosum and the abundance of axonal fascicles in the striatum were significantly reduced in cKO brains. At P90, the axonal phenotypes of cKO corpus callosum and striatum remain similar to that of P16/21 cKO brains. These data suggest the loss of NMNAT2 in post-mitotic glutamatergic neurons results in an age-dependent loss of long-range axons.

**Fig 2.**
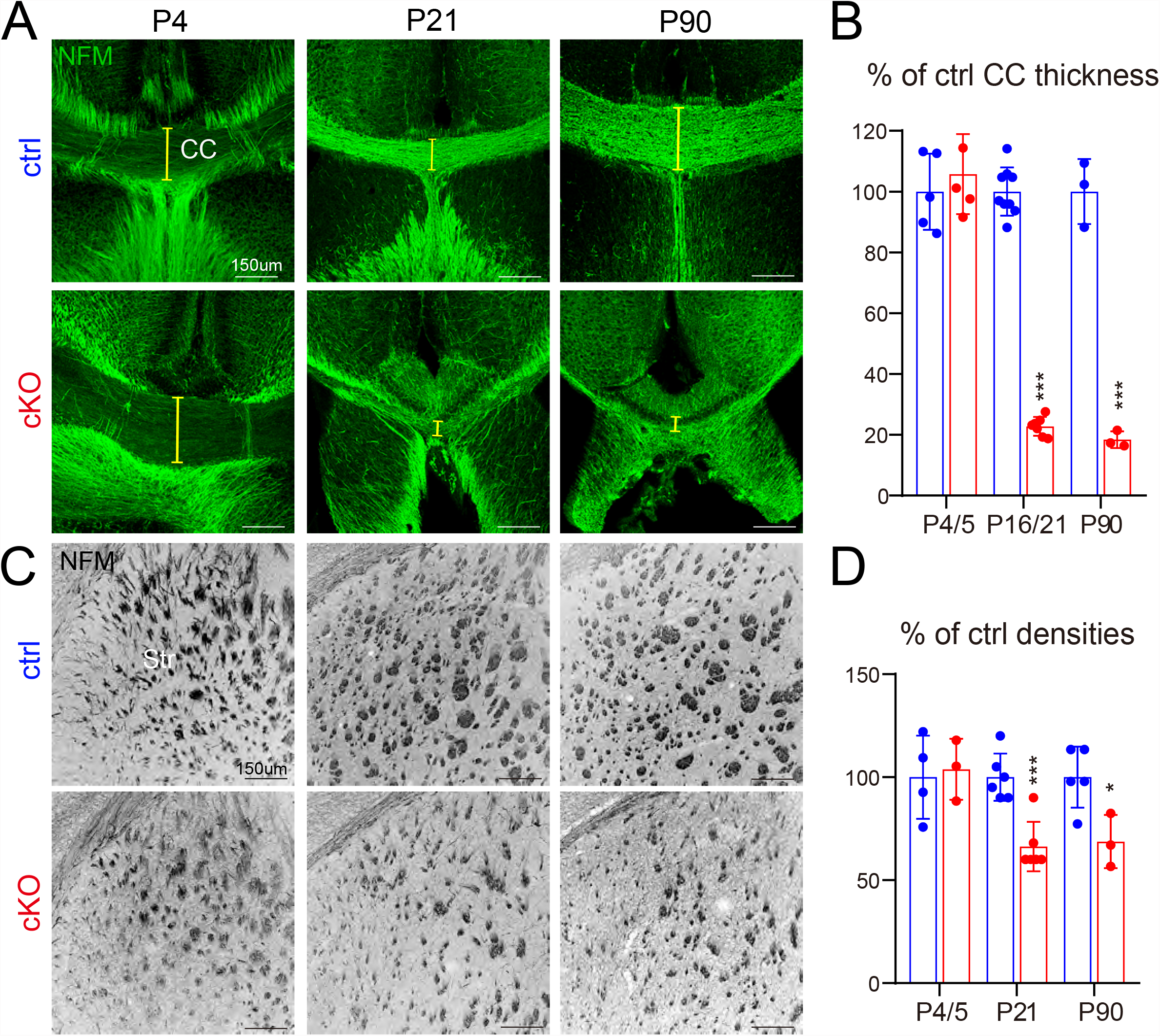
NMNAT2 cKO mice exhibit progressive loss of long-range cortical axons. (**A**) Representative images show axonal tracts revealed by NFM (neural filament M) immunoreactivity through the corpus callosum (CC) in ctrl and cKO at P4, P21, and P90. Yellow brackets mark where CC thickness was measured. (**B**) Summary for normalized CC thickness (P4/5 Ctrl n=5, cKO n=5; P16/21 Ctrl n=9, cKO n=7; P90 Ctrl n=3, cKO n=3). (**C**) Representative NF-M staining images showing axonal tracts through ctrl and cKO striatum (Str). (**D**) Summary for normalized NFM pixel densities in the striatum of ctrl and cKO mice (P4/5 Ctrl n=4, cKO n=3; P16/21 Ctrl n=6, cKO n=6; P90 Ctrl n=5, cKO n=3). **, p<0.01; ***, p<0.001 by Student’s t-test.

### Increased Inflammatory response and gliosis in NMNAT2 cKO mouse

Enlarged ventricles, reduced hippocampi and cortex, and age-dependent progression of axonal loss in cortical glutamatergic NMNAT2 KO brains are common features observed in neurodegenerative conditions (Coupe et al., 2019; Leyns et al., 2017). Inflammatory responses are often associated with neurodegenerative diseases such as AD, PD, and ALS (Chitnis and Weiner, 2017). To examine if NMNAT2 loss in cortical neurons triggered inflammatory responses, we globally evaluated the distribution of reactive astrocytes that are positive for GFAP (Glial fibrillary acidic protein; (Brahmachari et al., 2006; Chen et al., 1993; Eng and Ghirnikar, 1994; Eng et al., 1971)) and microglia that are positive for Iba1, a microglia-specific calcium-binding protein (Ito et al., 1998; Ohsawa et al., 2000) using immunostaining of coronal brain sections prepared from P5, P16/21, and P90 cKO and littermate control brains.

In general, we found fewer GFAP+ astrocytes in the cortex compared to hippocampi and striatum in control and cKO brains across the ages examined. No obvious increase of GFAP+ astrocytes was detected in cKO cortex, where the majority of neuronal somata with NMNAT2 deletion were located (Fig. 3A). However, in cKO brains, a substantial increase of GFAP+ immunoreactivity was observed in regions where long-range axonal tracts emanating from cortical glutamatergic neurons are located, such as white matter, corpus callosum, and striatum. Different from cortex, control hippocampi contain a plethora of GFAP+ astrocytes particularly enriched in the stratum lacunosum moleculare region located between CA1 and dentate gyrus (Fig. 3B). However, the area of GFAP+ signals was further increased in the whole hippocampi area in cKO brains. High magnification evaluation found that astrocytic morphologies reflected by GFAP signals varied in different brain regions as expected (Fig. 3C; (Escartin et al., 2021)). In addition, the morphologies of cKO-GFAP+ astrocytes were rather different from control-GFAP+ astrocytes from the corresponding brain regions. The hypertrophy patterns of cKO-GFAP+ astrocytes likely reflect changes in the astrocytic cytoskeleton (Escartin et al., 2021). Quantitative analysis found a significant increase of striatal GFAP+ signal already by P5, with the increase peaking at P16/21 and remaining significantly increased at P90 (Fig. 3D). However, in the hippocampi, GFAP+ signal densities were only significantly different between cKO and control brains at P16/21 but not at P5 or P90.

**Fig 3.**
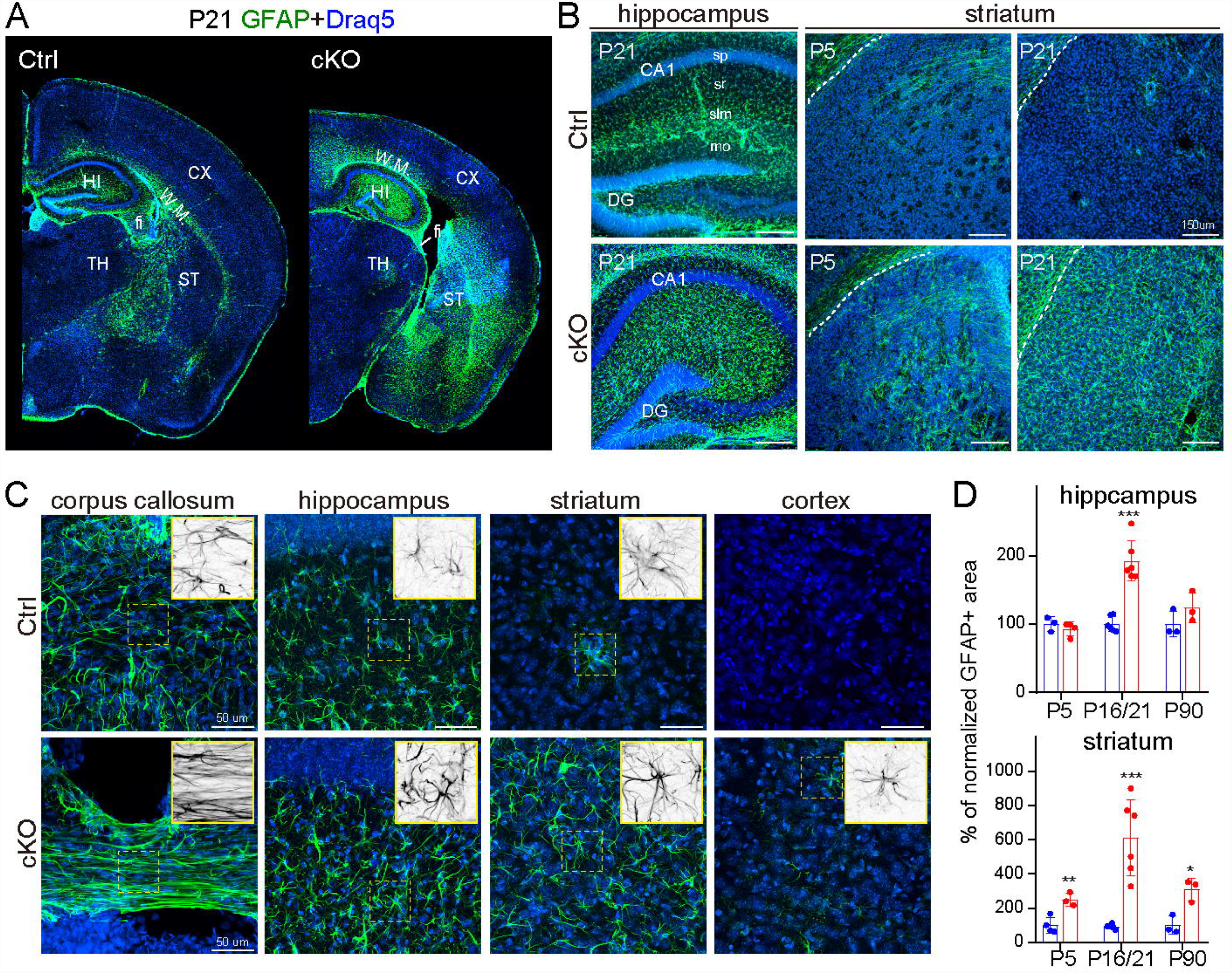
Deleting NMNAT2 from cortical glutamatergic neurons significantly increases reactive astrocytes around cortical axon tracts. (**A-C**) Example images of GFAP (green) and Draq5 (blue) staining in ctrl and cKO brains at P5 and P21. (**A**) Stitched overview images show increased GFAP immunoreactivity in brain regions where NMNAT2 KO long-range cortical axons pass through. (**B**) Representative GFAP staining images taken from the hippocampus and striatum. (**C**) High magnification images with inserts showing the morphologies of GFAP (black) positive astrocytes. (**D**) Summary for normalized GFAP+ area (P4/5 Ctrl n=3, cKO n=3; P16/21 Ctrl n=6, cKO n=6; P90 Ctrl n=3, cKO n=3). *, p<0.05; **, p<0.01; ***, p<0.001 by Student’s t-test. Abbreviations, DG, dentate gyrus; fi, fimbria; mo, moleculare layer in DG; slm, stratum lacunosum moleculare in CA1; sp, stratum pyramidale in CA1; sr, stratum radiatum in CA1; TH, thalamus; W.M., white matter.

Next, we examined the distributions and densities of Iba1-positive microglia in similar brain regions between control and NMNAT2 cKO mice at three ages (Fig. 4A). High magnification evaluations reveal different Iba1+ microglia morphologies in various brain regions as expected from previous studies (Fig. 4B; (Streit et al., 1999)). Interestingly, Iba1+ microglia in cKO brains often exhibit reduced ramifications (e.g. increase soma size and reduced process numbers), a sign of activated microglia. Similar to GFAP+ astrocytes, we found significant increases of Iba1+ microglia in cKO striatum from P5, further augmented at P16/21 but decreasing quite a bit by P90. However, there were significantly more Iba1+ microglia in cKO compared to control at all ages examined (Fig 4C). In the hippocampi, Iba1+ microglia densities were significantly higher than controls at P21 and P90. Taken together, our data suggest that deleting neuronal NMNAT2 results in neuroinflammation near long-range axons projecting from cortical glutamatergic neurons.

**Fig 4.**
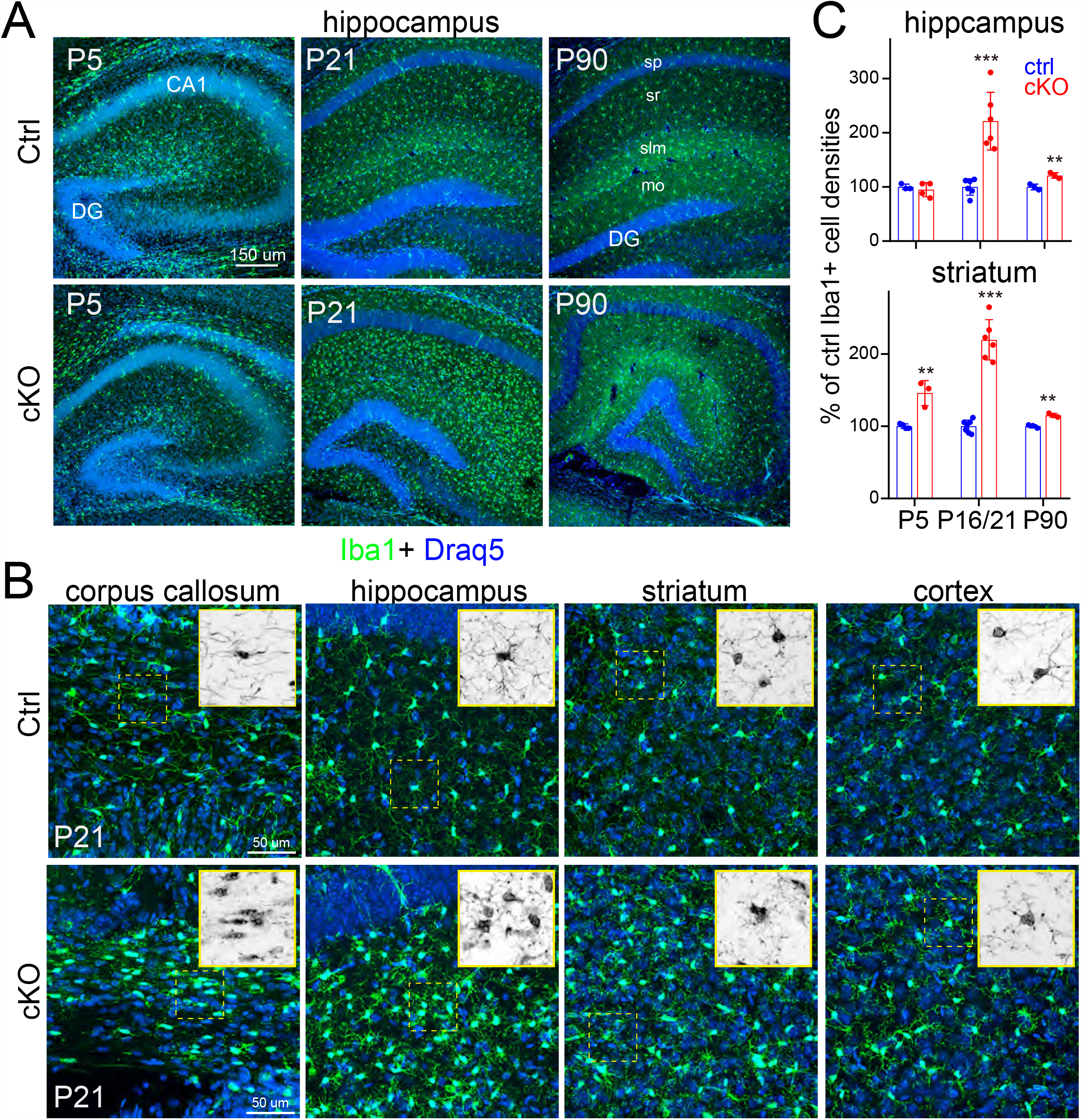
Deleting NMNAT2 from cortical glutamatergic neurons significantly increases the number of microglia. (**A-B**) Examples of images of Iba1 (green) and Draq5 (blue) staining taken from the hippocampus (**A**) and other brain regions of ctrl and cKO brains at P5, P21 and P90. (**B**) High magnification with inserts showing the morphologies of Iba1 positive (black) microglia. (**C**) Summary for normalized Iba+ cell densities in the hippocampus and striatum of ctrl and cKO brains (P4/5 Ctrl n=4, cKO n=3; P16/21 Ctrl n=7, cKO n=6; P90 Ctrl n=3, cKO n=3) **, *** indicate p<0.01, p<0.001 respectively by Student’s t-test.

### Unbiased transcriptomic analysis reveals upregulation of energy metabolism and downregulation of axonal outgrowth upon NMNAT2 loss in cortical neurons

The evidence that axonal loss does not occur in cKO corpus callosum until P16/21 demonstrates NMNAT2’s role in maintaining axonal health after initial outgrowth. To further understand the mechanism of the neuron-intrinsic axonal maintenance function mediated by NMNAT2, we adopted *in vitro* neuron-enriched primary culture model to examine the acute transcriptomic changes caused by NMNAT2 loss after Days-in-Vitro 9 (DIV9), the stage when cultured cortical neurons have accomplished their initial axonal outgrowth (Lesuisse and Martin, 2002). Specifically, we employed a Cre-loxP approach by lentiviral transduction to express Cre recombinase and delete NMNAT2 in cortical neurons cultured from NMNAT2^f/f^ embryos. By DIV12, 72 hours post lentiviral transduction, NMNAT2 expression is reduced by ∼88% in cKO neurons, compared to control neurons transduced by lentivirus expressing copGFP. Next, RNA-seq analyses were conducted with mRNA extracted from DIV12 control and cKO cortical neurons. Acute NMNAT2 loss resulted in significant upregulation of 1066 genes and downregulation of 1360 genes (FDR<0.05), which were further analyzed by web based Enrichr for gene ontology (GO) enrichment analysis with 2021 Biological Process terms (Xie et al., 2021). From upregulated genes, the top 10 over-represented Biological Processes identified by GO were enriched in bioenergetic functions, involving mitochondrial respiration, pyruvate metabolism and glycolysis, as well as translational regulation and autophagy (Fig. 5A). The pathways in mitochondrial metabolism with significantly upregulated genes highlighted, including fatty acid β oxidation, TCA cycle, pyruvate metabolism and oxidative phosphorylation, are delineated in Fig. 5B. From downregulated genes, GO analysis reveals top 10 Biological Processes in neurite development, involving axon guidance, axonogenesis and neuron projection morphogenesis, as well as extracellular organization, transcriptional regulation and calcium ion transport (Fig. 5A). The examples of significantly dampened axon guidance pathways in cKO neurons are illustrated in Fig. 5C. Of note, axonal terminals have shifted from their axonal outgrowth program towards synaptogenesis by DIV12 (Lesuisse and Martin, 2002), when a series of canonical axonal guidance molecules alternatively exert synaptogenic activity (Henderson and Dalva, 2018; Zou, 2020). We uncovered that many axon guidance signaling pathways essential for synaptogenesis, including Integrins, Ephrin receptors, Celsr3, Frizzeled3, Netrin 2G, DCC, Neurexins, Slit1/3, Robo2, Semaphorins and Plexin receptors, were downregulated (Fig. 5C). In addition, the activity-dependent immediate early genes, Egr3 and Egr4 were downregulated, as well as the neurotrophic factor Vgf, recently identified as a key regulator of Alzheimer’s disease (Beckmann et al., 2020). Interestingly, NMNAT2 loss also leads to the substantial elevation of Prion expression, whose misfolding trigger proteostress detrimental to neuronal health, and the reduction of Lrp1, which controls intracellular trafficking and extracellular spreading of Prion (Parkyn et al., 2008). Taken together, the transcriptomic analysis unbiasedly reveals the immediate consequence of NMNAT2 loss in perturbating the bioenergetic homeostasis, the axon guidance program, the activity-dependent synaptogenesis program and the proteostasis.

**Fig 5.**
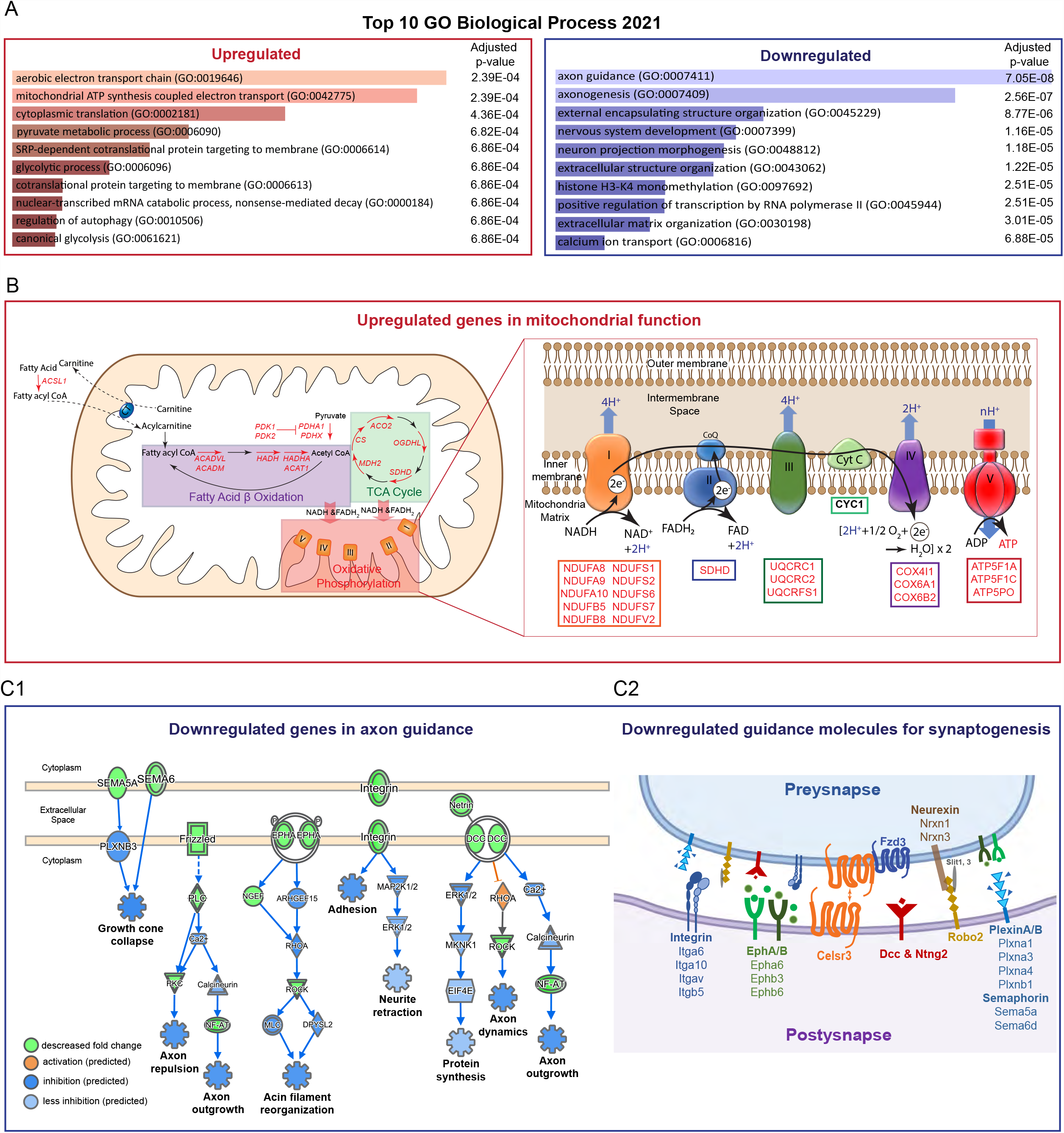
Transcriptomic profile of cortical neurons after acute deletion of NMNAT2 after neurite outgrowth. (**A**) Top 10 over-represented gene ontology terms of “2021 Biological process” ranked by *p*-value. (**B**) Upregulated genes (highlighted in red) in pathways of mitochondrial metabolism, including fatty acids β oxidation, TCA cycle, pyruvate metabolism, and oxidative phosphorylation. (**C1**) Downregulated genes (labeled in light green) in axon guidance pathways. (**C2**) Downregulated axon guidance molecules are genes expressed in both pre and post-synapse and are essential for synaptogenesis.

### Deleting NMNAT2 in cortical glutamatergic neurons results in the loss of the whisker-representation somatosensory map in the primary somatosensory (S1) cortex

Given the massively perturbed activity-dependent synaptogenesis process revealed by transcriptomic profiles in cortical neurons *in vitro*, we wanted to further investigate the impact of NMNAT2 loss on cortical circuit formation *in vivo*. The discrete anatomical organization of whisker-barrel map in the cortical layer 4 of S1 cortex (Steffen and Van der Loos, 1980; Woolsey et al., 1975; Woolsey and Van der Loos, 1970) and the clear one-to-one relationship of one whisker to one barrel have made this system a popular model to study neural activity-dependent process in cortical circuit formation (Erzurumlu and Gaspar, 2012; Fox, 2008; Wu et al., 2011). Each whisker-representing module is composed of thalamic axon clusters transmitting sensory information from a single whisker and a ring of layer IV neurons receiving thalamic inputs by projecting their dendrites toward the center, called barrels. The match between thalamocortical axons and corresponding cortical neurons for particular whiskers depends on glutamate transmission (Wu et al., 2011). To reveal the distributions of thalamocortical synapses in S1 cortex, VGluT2 and PSD95 double immunostaining together with the nuclear stain Draq5 was conducted in comparable brain sections containing S1 cortex from both P16/21 ctrl and cKO brains (Fig. 6A). VGluT2 immunoreactivity reveals thalamic axons originated from neurons in the thalamus with wild-type level NMNAT2 expression (Nahmani and Erisir, 2005), while PSD95 marks the post-synaptic compartments of cortical neurons that have had NMNAT2 eliminated. While the evident whisker-related patterns of thalamocortical synapses as well as the barrel cytoarchitecture were observed in the control S1 cortex, continuous distributions of VGluT2 and PSD95 signals without segregation were observed in cKO S1 cortex. This finding suggests that NMNAT2 loss in the cortical glutamatergic neurons disrupts the formation of whisker-related patterns, by interfering the barrel cytoarchitecture formation cell-autonomously, and by preventing the segregations of thalamic axons with wild-type levels of NMNAT2 expression in a non-autonomous manner. This observation suggests that NMNAT2 in cortical glutamatergic neurons participates in the glutamate-transmission dependent process of coordinating the establishment of whisker-representations between the presynaptic thalamocortical axons and the postsynaptic dendrites from cortical neurons.

**Fig 6.**
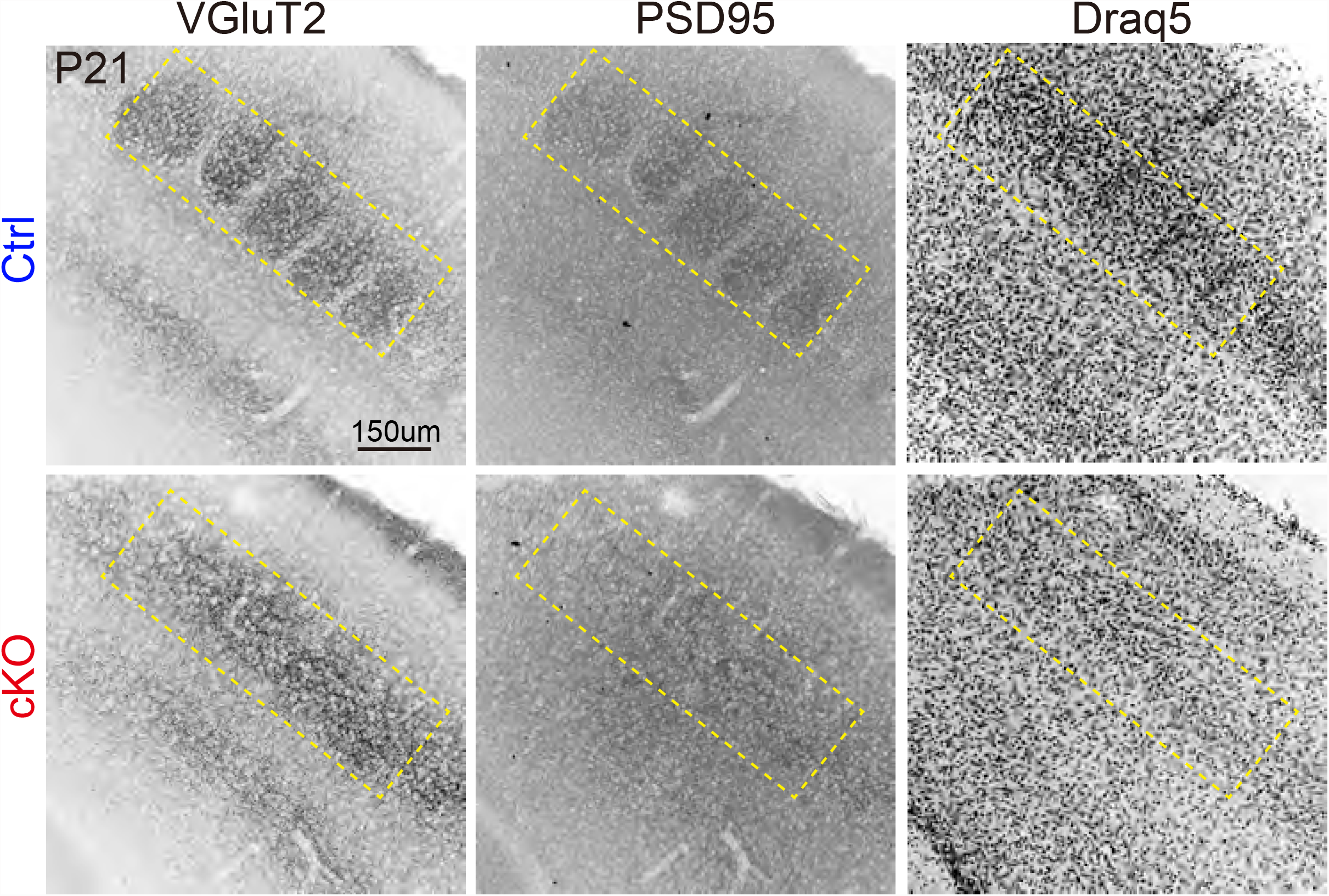
Cortical NMNAT2 deletions results in defective subcortical innervations into the S1 cortex and hippocampal area. Representative images with color inverted show VGluT2, PSD95, and Draq5 staining in the S1 cortex of ctrl (n=5) and cKO (n=5) brains.

### Knocking out one allele of Sarm1 in NMNAT2 cKO mice rescues somatosensory map formation but not axonal loss and inflammation

Sarm1 (Sterile alpha and TIR motif containing1) is a NAD(P) glycohydrolase and a prominent executor of programmed axon death (Angeletti et al., 2022; Hopkins et al., 2021). Compelling evidence shows that an increase in NMN to NAD+ ratio after NMNAT2 loss activates Sarm1 to trigger axon degeneration (Figley et al., 2021). Deleting Sarm1 is neuroprotective against axon injury and increases neuronal survival (Conforti et al., 2014; Peters et al., 2021). NMNAT2 global KO mice die perinatally (Hicks et al., 2012), while Sarm1 deletion allows these mice to survive and stay overtly healthy for up to 24 months (Gilley et al., 2015). Despite comprehensive studies of protection by Sarm1 deletion on peripheral nerves in NMNAT2 global KO mice (Gilley et al., 2015), it remains unknown whether Sarm1 deletion rescues brain specific phenotypes caused by NMNAT2 loss. To determine whether reducing Sarm1 abundance reverses brain specific phenotypes caused by NMNAT2 loss, we crossed NMNAT2 cKO mice to transgenic mice carrying Sarm1-null alleles (abbreviated as S^null^) to generate NMNAT2 cKO in Sarm1-null heterozygous (cKO;S^null^/+) or homozygous background (cKO;S^null^/S^null^). Surprisingly, we found that deleting one allele of Sarm1 (cKO;S^null^/+) is sufficient to rescue the whisker-representation somatosensory map in S1 cortex that is ablated by NMNAT2 loss (Fig. 7). Conversely, at the gross level, the brains from cKO;S^null^/+ mice still exhibited aberrant anatomy similar to NMNAT2 cKO mice as described earlier (Fig. 1E), such as reduced body weight, smaller hippocampi, enlarged ventricles and reduced thickness of cortex and corpus callosum (CC) (Sup-Fig. 3). Immunostaining of NF-M also reveal significant axonal loss in both CC (Fig. 8A, B) and striatum (Fig. 8C, D). Inflammatory responses reflected by a dramatic increase in the density of GFAP+ astrocytes and iba1+ microglia were transiently evident at P21 in cKO;S^null^/+ brains (Fig. 9), however this evidence of inflammation ultimately disappear by P90. When two alleles of Sarm1 are removed, none of the NMNAT2 cKO phenotypes were detectable, reinforcing the observation that the complete Sarm1 loss of function antagonizes the brain specific phenotypes caused by NMNAT2 loss.

**Fig 7.**
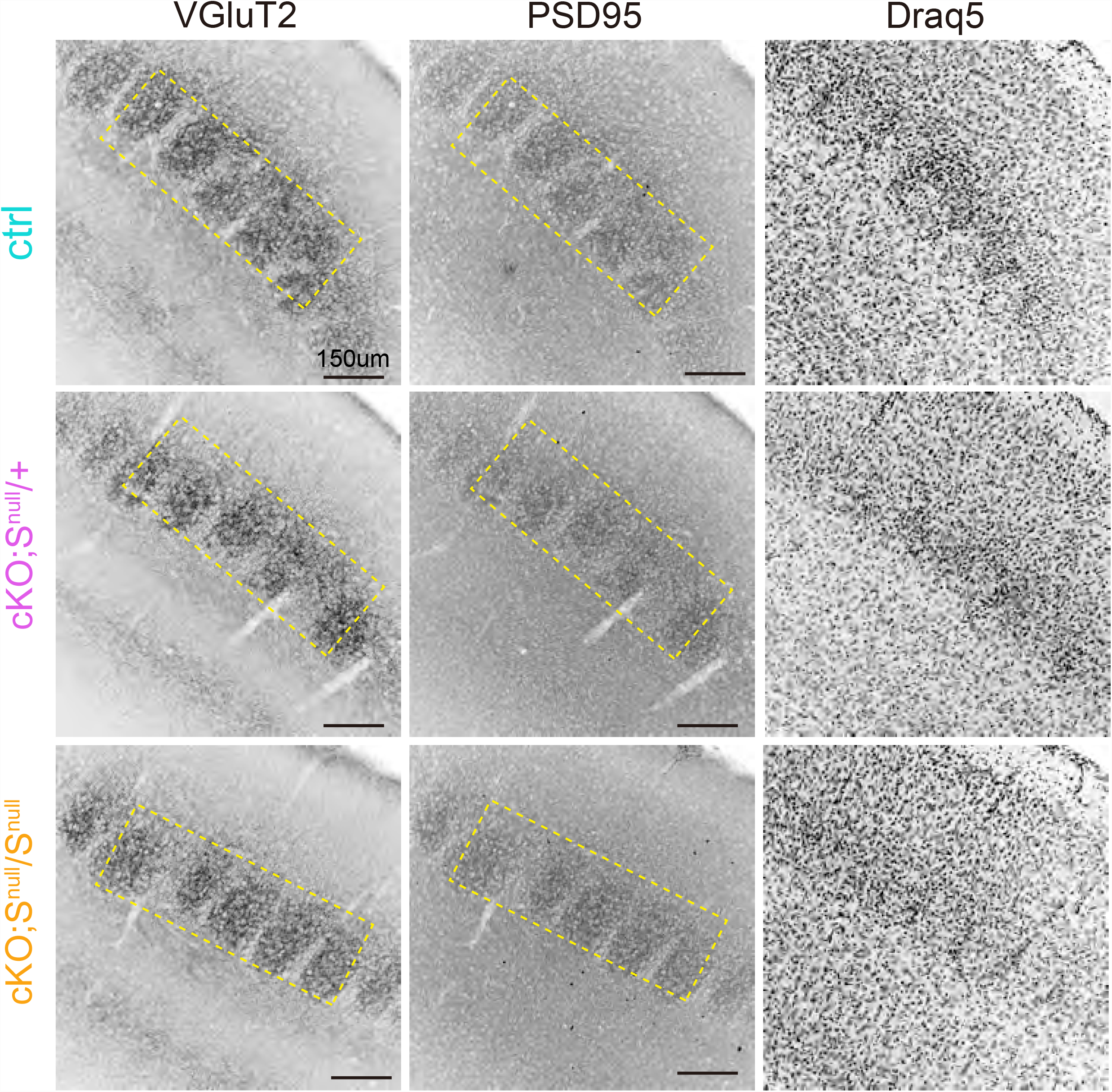
Sarm1 reduction rescues the whisker-barrel map deficits in NMNAT2 cKO mice. Inverted color images show the VGluT2, PSD95 or Draq5 staining in the S1 cortex of ctrl (n=10), NMNAT2 cKO;Sarm1^null^/+ (cKO;S^null^/+; n=14), NMNAT2 cKO;Sarm1^null /null^ (cKO;S^null^/S^null^; n=5) brains.

**Fig 8.**
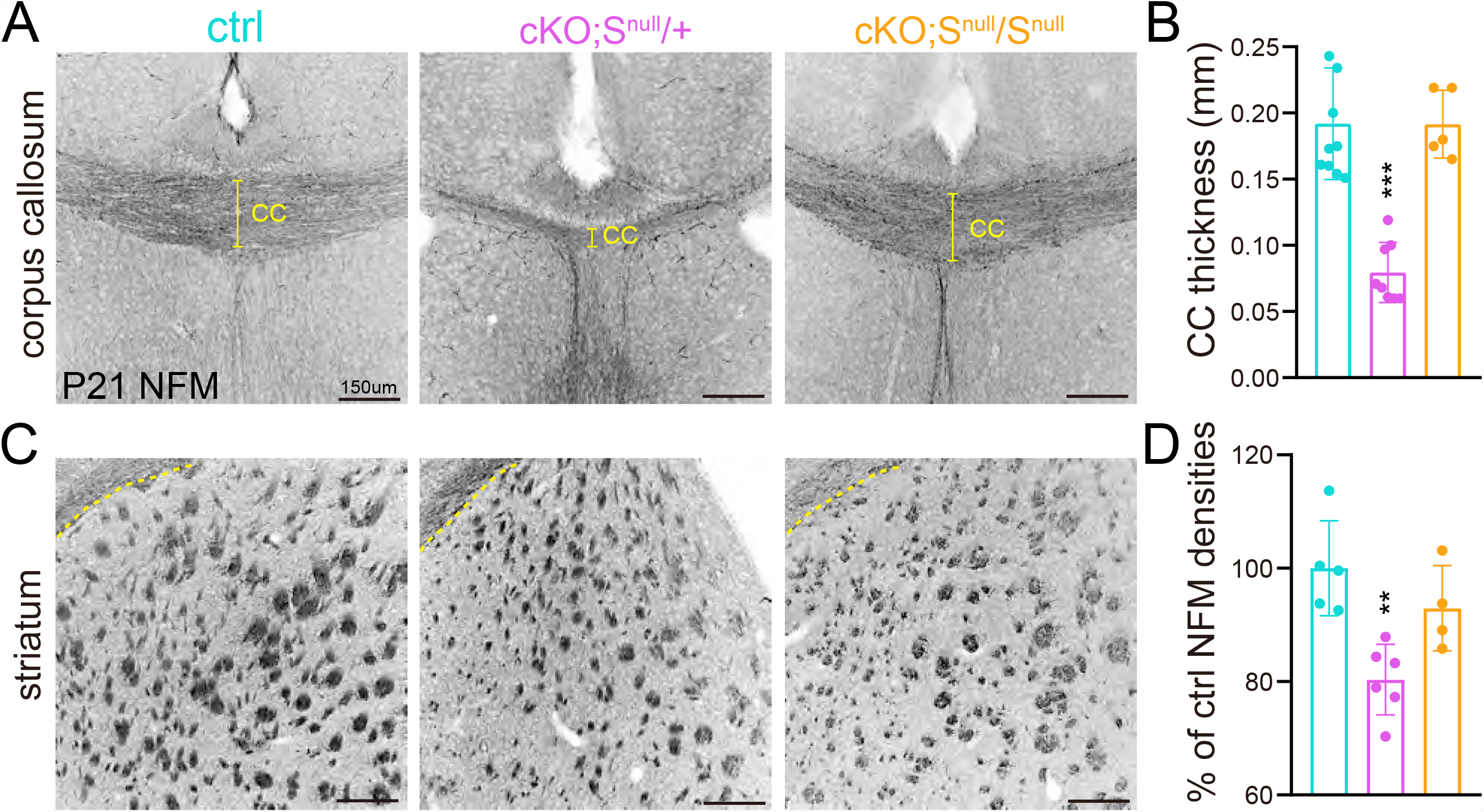
Reducing Sarm1 abundance ameliorates axonal phenotypes caused by NMNAT2 deletion. (**A, C**) Representative NF-M staining (black) images show axons passing through the corpus callosum (**A**) and striatum (**C**) in P21 ctrl, cKO;S^null^/+, cKO;S^null^/S^null^ brains. (**B, D**) Summaries for normalized CC thickness (ctrl, n=10; cKO;S^null^/+, n=8; cKO;S^null^/S^null^, n=5) and normalized NFM pixel densities in the medial region along rostral-caudal axis of the striatum (ctrl, n=5; cKO;S^null^/+, n=6; cKO;S^null^/S^null^, n=5). **, *** indicate p<0.01 or p<0.001.

**Fig 9.**
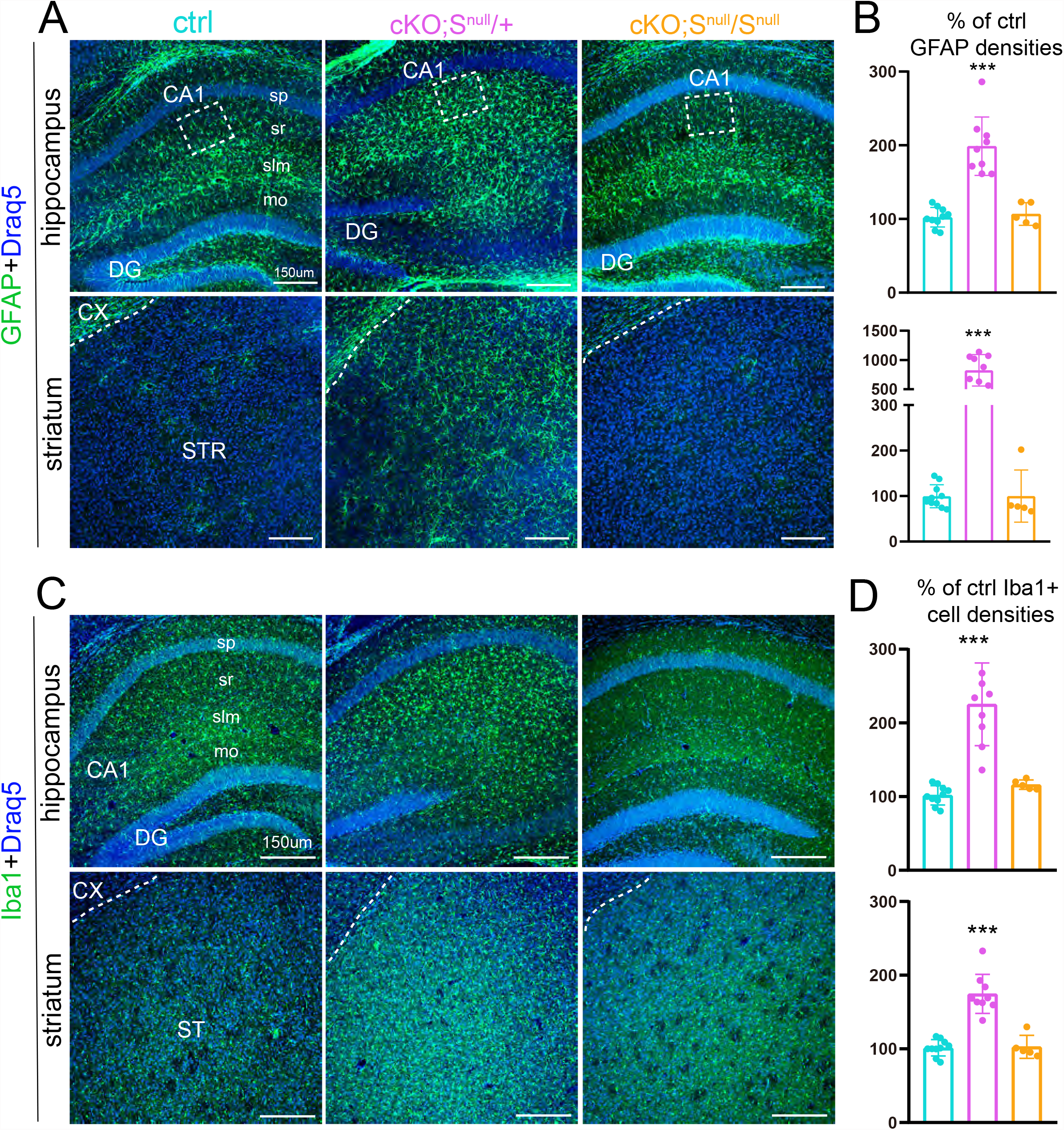
Reducing Sarm1 abundance attenuates the inflammatory responses caused by NMNAT2 deletion. (**A**) Example GFAP staining images taken from the hippocampus and striatum of P21 ctrl, cKO;S^null^/+, cKO;S^null^/S^null^ brains. (**B**) Summary for normalized GFAP densities. (**C**) Example Iba1 staining images taken from the hippocampus and striatum of P21 ctrl, cKO;S^null^/+, cKO;S^null^/S^null^ brains. (**D**) Summary for normalized Iba1 cell densities (P21 ctrl n=10; cKO;S^null^/+, n=9; cKO;S^null^/S^null^, n=5). ***, p<0.001.

## Discussion

Here we show that NMNAT2 in cortical glutamatergic neurons is required for proper cortical development including cortical somatosensory map formation as well as for maintaining axonal health and integrity. Despite the specific deletion of NMNAT2 in post-mitotic glutamatergic neurons from a late embryonic age, dramatic phenotypes were observed even at the global level, including reduced survival, reduced body weights, and smaller brain sizes. Age-dependent reduction of long-range axonal tracts originating from cortical glutamatergic neurons and a massive increase in inflammatory responses in cKO brains from weaning onwards suggest that NMNAT2 is required immediately after the formation of axonal tracts to maintain healthy axons. The enrichment of reactive astrocytes and activated microglia in brain regions with traversing long-range cortical axons, highlight the vulnerability of long-range axonal projections upon NMNAT2 loss. Unbiased transcriptomic studies suggest that NMNAT2 loss in cortical neurons attenuates biological processes related to axonal outgrowth and synaptogenesis while increasing mitochondria function. Compelling evidence show that NMNAT2 loss activates Sarm1 to trigger axon degeneration and loss of Sarm1 protect axons from degeneration in the peripheral nervous system (27, 28). Here we found that completely deleting Sarm1 rescues all the observed CNS phenotypes in NMNAT2 cKO mice. But if only one genomic copy of Sarm1 is inactivated, NMNAT2 cKO brains still exhibit axonopathy and inflammatory responses in corpus callosum and striatum, however whisker-barrel map formation is largely intact. Taken together, our study provides the first evidence for the roles of NMNAT2 in both the development and maintenance of cortical neural circuits. The restoration by Sarm1 loss-of-function, a NAD(P) glycohydrolase, emphasizes the importance of the NAD^+^ synthesis activity provided by NMNAT2 for normal CNS circuit development and maintenance.

### NMNAT2 deletion from cortical glutamatergic neurons results in neurodegenerative-like phenotypes from very early postnatal ages

Several neurodegenerative-like phenotypes were observed in NMNAT2 cKO mice: (1) ataxia, hindlimb clasping (Fig. 1), and forelimb circling (data not shown) motor behavior phenotypes; (2) enlarged ventricles; (3) reduced cortical thickness and shrunken hippocampi; (3) age-dependent reduction in axonal tracts and the thickness of the corpus callosum and anterior commissure (Figs 1-2); and (4) massive increase in the numbers of reactive astrocytes and activated microglia (Fig. 3). However, the early onset of these dramatic phenotypes, at the age of weaning suggests that NMNAT2 plays a key role in brain development as well as in neuronal maintenance. Indeed, NMNAT2 is expressed in post-mitotic glutamatergic neurons from embryonic stages and the whisker-barrel map failed to form in cKO S1 cortices. Previous experiments using cortical explant cultures found reduced axonal outgrowth from NMNAT2 KO cortex (Gilley et al., 2013a). We found normal thickness of corpus callosum in cKO brains during the first postnatal week (Fig. 2), suggesting early formation of axonal tracts is preserved. Our preliminary analysis of P7-P90 mosaic mice lacking NMNAT2 in only a few cortical neurons embedded in an overall wildtype background found NMNAT2 KO axons originating from cortical layer2/3 glutamatergic neurons cross the corpus callosum and reach their cortical target area in the contralateral hemisphere (data not shown). This suggests initial axon outgrowth from cortical neurons to their target zone does not require NMNAT2. However, NMNAT2 KO axons in the wildtype environment exhibit reduced numbers of axonal arbors in their target zones compared to control axons (data not shown). *In vitro* cortical neuronal culture studies found normal neurite densities formed by KO cortical neurons (data not shown). Taken together, our findings suggest that NMNAT2 is not required for the initial axonal outgrowth but is likely to be important for maximizing axonal arborization.

### Unbiased transcriptomic analysis reveals the impacts of NMNAT2 deletion on mitochondria function and axon formation/synaptogenesis

Our unbiased transcriptomic analysis designed to identify differentially expressed genes after ∼2 days of NMNAT2 reduction found the upregulation of several pathways in mitochondrial metabolism and downregulation of pathways related to axonal function. In our study, we deleted NMNAT2 after most cortical neuron neurite outgrowth had been completed and during the initiation of synaptogenesis. The majority of the cells (∼85%) in our cortical cultures are glutamatergic neurons with low numbers of astrocytes and inhibitory neurons (data not shown). Thus, the transcriptomic changes we detected are likely to reflect the direct impacts of NMNAT2 loss on glutamatergic neurons. Numerous genes that are upregulated upon NMNAT2 loss are involved in fatty acid β oxidation, TCA cycle, and oxidative phosphorylation (Fig. 5). It remains to be determined whether these transcriptional changes result in enhanced mitochondrial function and subsequently increase oxidative stress and trigger neuroinflammation *in vivo*. Many of the genes downregulated after NMNAT2 loss were annotated to axonal outgrowth and axonogenesis biological processes. However, recent work suggests that many classical axonal guidance molecules including the Wnt/Frizzle, Ephrin/Eph, Slit/Robo, Semaphorin/Plexin and Integrin families also play critical roles in synaptogenesis and maintenance (Henderson and Dalva, 2018; Zou, 2020). Frizzled3 is enriched in synaptic vesicles and on the plasma membrane of presynaptic boutons whereas Celsr3 is present on the plasma membranes of both pre- and postsynaptic neurons and promotes synapse formation (Teo and Salinas, 2021; Thakar et al., 2017; Zou, 2020).

The whisker-related organization of both pre- and post-synaptic components of thalamocortical connections fail to develop in NMNAT2 cKO S1 cortex, despite the deletion of NMNAT2 only from cortical neurons. The “barrelless” phenotype of glutamatergic-NMNAT2 cKO mice, together with the down-regulation of proteins involved in synaptogenesis, suggest an important role of NMNAT2 in neural activity-dependent synaptogenesis critical for cortical circuit formation. Interestingly, the normal whisker map in cKO;S/+ mice suggests ∼50% reduction of Sarm1 is sufficient to prevent the impact of NMNAT2 loss on this activity-dependent process.

## Funding and Acknowledgments

This work was supported by the NIH-NINDS R01NS086794 (HCL). We thank Drs. Ken Mackie and Jui-Yen Huang for helpful comments. We also thank Scott Barton for technical assistance. Confocal images were taken in the Light Microscopy Imaging Center at Indiana University Bloomington (NIH1S10OD024988). The sequencing was performed in the Center for Medical Genomics (CMG) at Indiana University School of Medicine, a sequencing core facility of the Indiana Clinical and Translational Sciences Institute. The *in situ* data presented was supported in part by the RNA In Situ Hybridization Core at Baylor College of Medicine supported by NIH IDDRC grant U54HD083092 from the Eunice Kennedy Shriver National Institute of Child Health & Human Development. J.G. was funded by BBSRC/AstraZeneca Industrial Partnership Award BB/S009582/1 and M.P.C. was funded by the John and Lucille van Geest Foundation.

## Supplementary Figures

**Sup Fig. 1.**
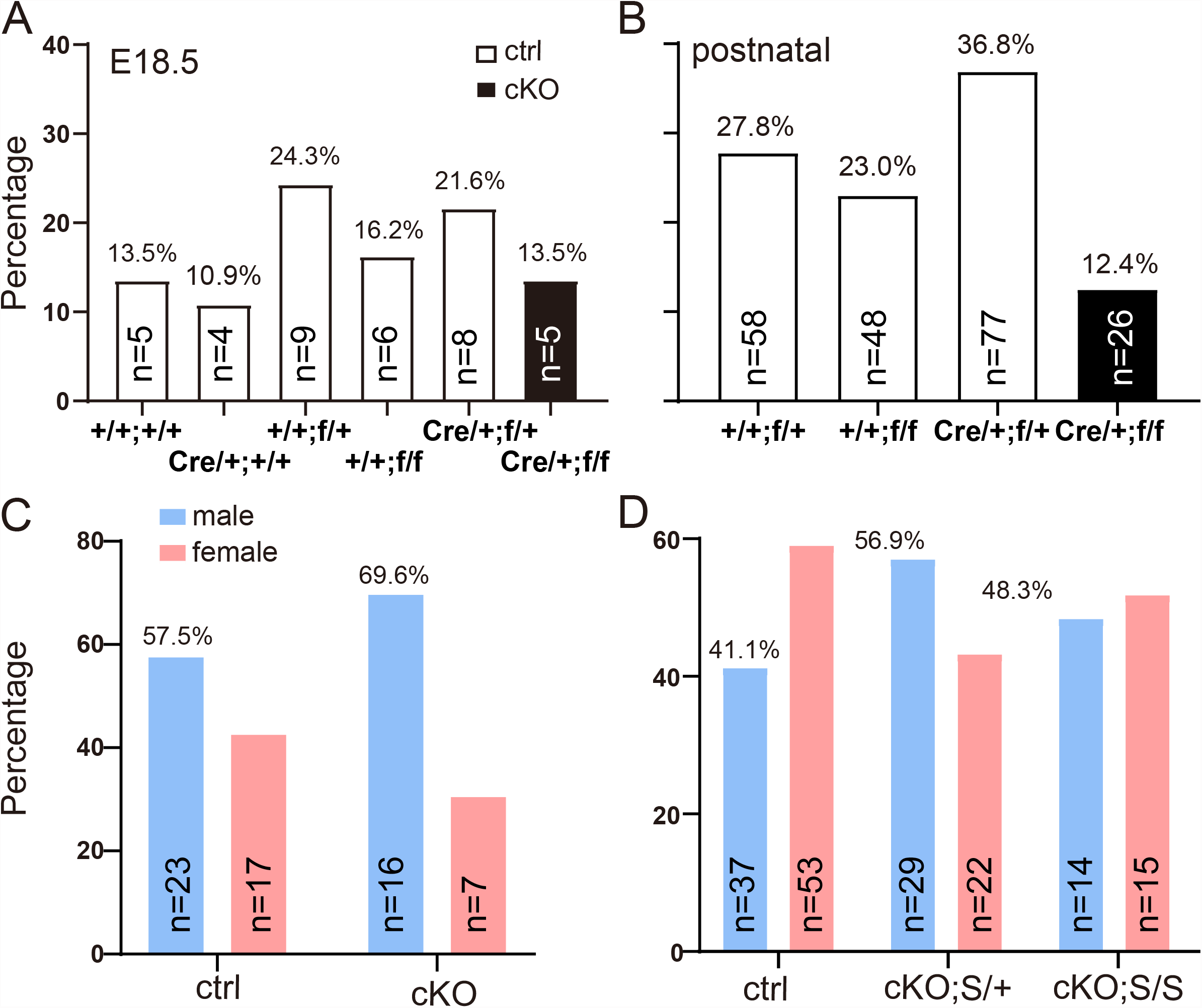
Survival rate and gender ratio. (**A**) Genotype percentage of prenatal (E18.5) NMNAT2 cKO mice. Parents genotype: NMNAT2^F/+^ Cre, and NMNAT2^F/+^. The offspring genotypes: NMNAT2 ^+/+^, NMNAT2 ^+/+^ Cre, NMNAT2 ^F/+^, NMNAT2 ^F/+^ Cre, and NMNAT2 ^F/F^, were clustered as Ctrl; NMNAT2 ^F/F^ Cre as cKO. Total N number = 37. (**B**) Genotype percentage of postnatal NMNAT2 cKO mice. Parents genotype: NMNAT2^F/+^ Cre, and NMNAT2^F/F^. The offspring genotypes: NMNAT2 ^F/+^, NMNAT2 ^F/+^ Cre, and NMNAT2 ^F/F^, were clustered as Ctrl; NMNAT2 ^F/F^ Cre as cKO. Total N number = 209. (**C**) Gender ratio of postnatal Ctrl and NMNAT2 cKO mice. Total Ctrl N = 40; cKO N = 23. (**D**) Gender ratio of postnatal Ctrl, cKO Sarm1^+/-^ (cKO;S/+) and cKO Sarm1^-/-^ (cKO;S/S) mice. Ctrl, n= 90; cKO;S/+, n=51; cKO;S/S, n= 29.

**Sup Fig. 2.**
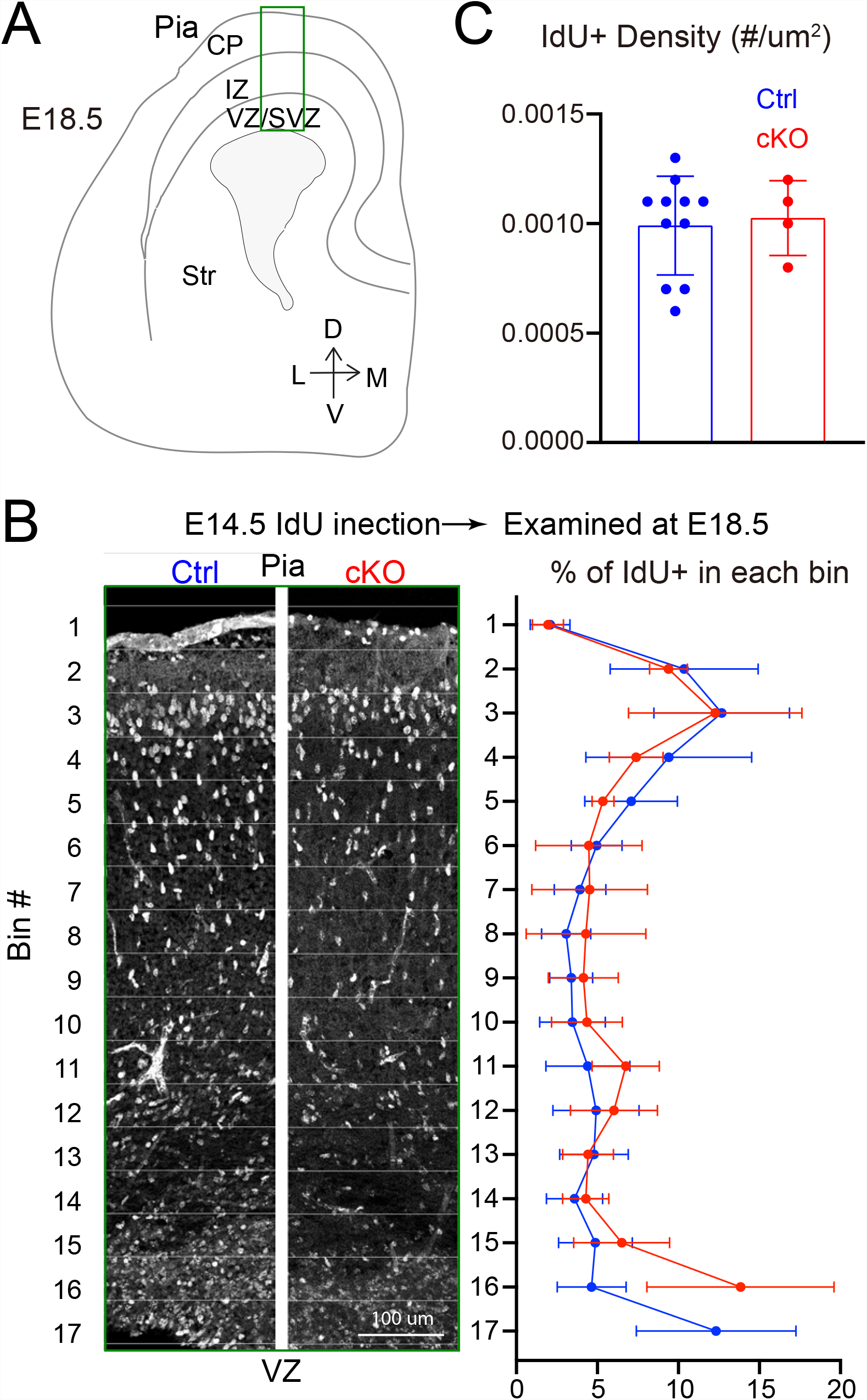
Deleting NMNAT2 from post-mitotic glutamatergic neurons does not cause obvious phenotypes during embryonic development. (**A**) Cartoon revealing the different cortical layers and morphological hallmarks of E18.5 brain, with the highlighted box showing the different layers of cortical plate (CP) region quantified in B. (**B-C**) IdU was injected at E14.5 and embryos were harvested at E18.5 to examine cell proliferation and migration in ctrl and NMNAT cKO embryos. (**B**) Example IdU staining in E18.5 cortical plate following IdU incorporation at E12.5. Right panel shown summaries for the distributions of IdU+ cells throughout the cortical plate. Quantification of IdU+ cells within square bins drawn from the ventricle to the marginal zone. (**C**) Summary for IdU+ cell densities. Ctrl, n=11; cKO, n=4.

**Sup Fig. 3.**
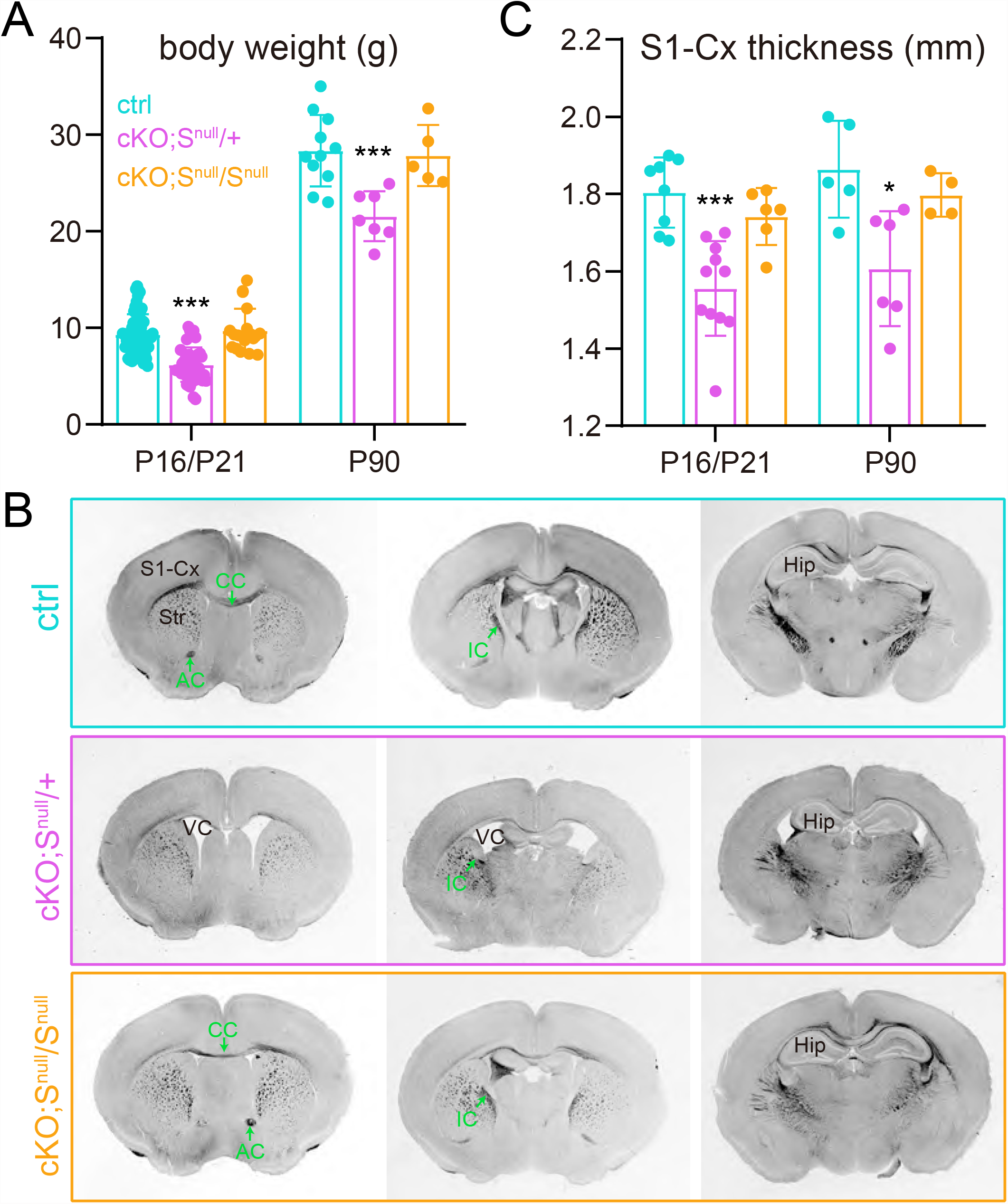
Sarm1-NexGTE global observation. (**A**) Body weight of ctrl, cKO;S^null^/+ and cKO;S^null^/S^null^ mice at P16/21 and P90. cKO;S^null^/+ mice show lighter body weight compared to ctrl and cKO;S^null^/S^null^ mice.(B) Brightfield coronal brain images for ctrl, cKO;S^null^/+ and cKO;S^null^/S^null^ mice. cKO;S^null^/+ show enlarged ventricles (VC), atrophied hippocampi (Hip), thinner corpus callosum (CC) and shrunk S1 cortex similar to cKO mice. However, cKO;S^null^/S^null^ show normal brain anatomy. (C) Quantification of the thickness of S1 cortex in ctrl, cKO;S^null^/+, and cKO;S^null^/S^null^ mice at P16/21 and P90. The cKO;S^null^/+ group showed significantly reduced cortical thickness at P21 and P90 compared to ctrl and cKO;S^null^/S^null^ mice.

## Notes

Conflict of Interest: None.

### Competing Interest Statement

The authors have declared no competing interest.

